# Depletion of TAX1BP1 amplifies innate immune responses during respiratory syncytial virus infection

**DOI:** 10.1101/2021.06.03.447014

**Authors:** Delphyne Descamps, Andressa Peres de Oliveira, Lorène Gonnin, Sarah Madrières, Jenna Fix, Carole Drajac, Quentin Marquant, Edwige Bouguyon, Vincent Pietralunga, Hidekatsu Iha, Armando Morais Ventura, Frédéric Tangy, Pierre-Olivier Vidalain, Jean-François Eléouët, Marie Galloux

## Abstract

Respiratory syncytial virus (RSV) is the main cause of acute respiratory infections in young children, and also has a major impact in the elderly and immunocompromised people. In the absence of vaccine or efficient treatment, a better understanding of RSV interactions with the host antiviral response during infection is needed. Previous studies revealed that cytoplasmic inclusion bodies (IBs) where viral replication and transcription occur could play a major role in the control of innate immunity during infection by recruiting cellular proteins involved in the host antiviral response. We recently showed that the morphogenesis of IBs relies on a liquid-liquid phase separation mechanism depending on the interaction between viral nucleoprotein (N) and phosphoprotein (P). These scaffold proteins are expected to play a central role in the recruitment of cellular proteins to IBs. Here, we performed a yeast two-hybrid screen using RSV N protein as a bait, and identified the cellular protein TAX1BP1 as a potential partner of N. This interaction was validated by pulldown and immunoprecipitation assays. We showed that TAX1BP1 suppression has only a limited impact on RSV infection in cell cultures. On the contrary, *in vivo* experiments showed that RSV replication is decreased in TAX1BP1^KO^ mice, whereas the production of inflammatory and antiviral cytokines is enhanced. *In vitro* infection of either wild-type or TAX1BP1^KO^ alveolar macrophages confirmed that the innate immune response to RSV infection is enhanced in the absence of TAX1BP1. Altogether, our results suggest that RSV could hijack TAX1BP1 to restrain the host immune response during infection.

**Importance:** Respiratory syncytial virus (RSV), which is the leading cause of lower respiratory tract illness in infants, still remains a medical problem in the absence of vaccine or efficient treatment. This virus is also recognized as a main pathogen in the elderly and immunocompromised people, and the occurrence of co-infections (with other respiratory viruses and bacteria) amplifies the risks of developing respiratory distress. In this context, a better understanding of the pathogenesis associated to viral respiratory infections, which depends on both viral replication and the host immune response, is needed. The present study reveals that the cellular protein TAX1BP1, which interacts with the RSV nucleoprotein N, participates in the control of the innate immune response during RSV infection, suggesting that N-TAX1BP1 interaction represents a new target for the development of antivirals.

## INTRODUCTION

Respiratory syncytial virus (RSV) is the main pathogen responsible for acute respiratory infections and bronchiolitis in children (1). Almost all children are infected by the age of two. A systemic multisite study on the cause of infant’s pneumonia in hospitalized children in Asia and Africa recently revealed that RSV is the main etiological agent of severe pneumonia, accounting for over 30% of infections (2). In the United States, RSV is estimated to be responsible for the hospitalization of 86,000 children per year, with a related cost of 394 million dollars (3). Furthermore, RSV infections in early childhood is recognized to later increase the susceptibility to chronic asthma (4, 5). Reinfections occur throughout life and, if healthy adults generally present symptoms of bad cold, RSV infections are associated with significant morbidity and mortality in the elderly and immunocompromised people (6-9). Indeed, RSV is estimated to cause over 17,000 deaths per year in the United States, 78% of which occur in adults over 65 years of age, and is responsible for 5% of total hospital admissions in the elderly (10). Although RSV has a major impact on human health and the economy, there is still no vaccine available. The development of vaccines has been hampered by the repercussions of a failed vaccine trial using a formalin-inactivated virus in the 1960s, which resulted in an exacerbation of the pathology upon infection and led to two deaths (11). The current standard of care consists of prophylactic treatment of at-risk infants with a monoclonal antibody (Palivizumab), but its use is limited by its moderate effectiveness and high cost (12).

The pathology associated to RSV infection results from both viral replication and the host’s immune response (13). RSV infection triggers an early immune response mediated by the production of type I interferons (IFN-I) which induces the transcription of IFN-stimulating genes (ISG) and the production of proinflammatory mediators (14-17). On the other hand, RSV has developed multiple strategies to hijack cellular pathways controlling the IFN-I and NF-κB (Nuclear Factor kappa B) pathway in order to blunt the host antiviral response (17-19). In particular, the two nonstructural viral proteins NS1 and NS2 are known to suppress IFN-I production and cell signaling during infection (20). Although IFN-I are major players in viral clearance and are essential to induce an appropriate immune response (21), they could also contribute to RSV pathogenesis with potentially different roles in infants and adults (17, 22-26). Indeed, high levels of IFN-I and inflammatory cytokines usually correlate with severity as this reflects the inability of the immune response to control the virus. It is thus essential to better characterize the complex interactions between RSV and the host immune response to decipher pathogenesis and design effective treatments.

RSV belongs to the *Mononegavirales* (*MNV*) order and the *Pneumoviridae* family (27). It is an enveloped virus with a non-segmented negative strand RNA genome containing 10 genes that encode 11 proteins. The two surface glycoproteins G and F are involved in the initial steps of infection, *i*.*e*. attachment and fusion with the cell membrane. The viral membrane, which also contains the small hydrophobic protein SH, is lined by the matrix protein M that is driving virus assembly. The genome is enwrapped by the nucleoprotein N, forming a helical nucleocapsid (28). The polymerase complex composed of the large polymerase (L) and its main cofactor the phosphoprotein P, is associated to this ribonucleoprotein complex (RNP) which serves as a template for viral transcription and replication (29). The viral transcription factor M2-1 is also present in the viral particle. After cell entry, RSV replicates in the cytoplasm of host cells within viro-induced spherical cytoplasmic granules called inclusion bodies (IBs). These structures are viral factories where all the viral proteins of the polymerase complex concentrate to perform the replication and transcription of the viral genome (30). These structures also play a role in viral escape from the innate immune system by limiting the recognition of viral RNAs by cytoplasmic pattern recognition receptors (PRRs) such as RIGI-I (Retinoic acid-Inducible Gene I) and MDA5 (Melanoma Differentiation-Associated gene 5). Once stimulated, these PRRs activate the transcription factors NF-κB and interferon regulatory factor 3 and 7 (IRF3/7) (31). The function of IBs in the modulation of the host innate immune response was further supported by a study showing that MDA5 interacts with the RSV-N protein. In addition, MDA5 and the downstream signaling molecule MAVS (Mitochondrial AntiViral Signaling both colocalize to IBs as soon as 12 hours post-infection, leading to downregulation of *IFNβ* mRNA expression (32). More recently, a study also revealed the sequestration of the NF-κB subunit p65 in RSV IBs (33). It is thus now admitted that the recruitment of cellular proteins into IBs participates not only to viral replication but is also involved in the control of cellular responses.

We recently showed that RSV IBs display hallmarks of liquid-liquid phase separation, and that the N and P proteins are at the core of the RSV IBs biogenesis (34). Their role as scaffold proteins suggest that N and P are directly involved in the partitioning of cellular proteins to IBs. However, their interactions with cellular factors are still poorly characterized. Here we report the identification of Tax1-binding protein 1 (TAX1BP1) as an interactor of RSV-N. TAX1BP1 was initially identified as a partner of the Tax protein from Human T-lymphotropic virus 1 (HTLV-1) (35). Since then, TAX1BP1 was shown to interact with viral proteins from Papillomaviruses (36), Measles virus (MeV) (37) and Mammarenaviruses (38). Among the described activity of TAX1BP1, this protein was involved in the negative regulation of NF-κB and IRF3 signaling by editing the ubiquitylation of its catalytic partner, the protein A20 (39, 40). We thus investigated the role of TAX1BP1 in both RSV replication and control the host antiviral response using *in vitro* and *in vivo* infection models. Altogether our results suggest that TAX1BP1 is hijacked by RSV to inhibit the host antiviral response.

## RESULTS

### Identification of TAX1BP1 interaction with the viral nucleoprotein N

To identify cellular interactors of the RSV-N protein, we first performed a yeast two-hybrid (Y2H) screen. Yeast cells were transformed with a vector encoding the RSV-N protein fused to GAL4 DNA binding domain (GAL-BD) in order to use it as bait in the Y2H system. Surprisingly, no yeast clones were obtained, suggesting that RSV-N is toxic. This could be due to the non-specific RNA-binding properties of N (41). We thus decided to use as a substitute the N protein harboring the K170A/R185A mutations that were previously shown to impair the interaction of N with RNA. This mutant is expressed as a monomeric RNA-free N, named N^mono^, which can mimic the natural N^0^ form (41). When yeast cells were transformed with a vector encoding N^mono^ fused to GAL4-BD, growing colonies were finally obtained on selective medium as expected. Yeast cells expressing N^mono^ were then mated with yeast cells transformed with a human spleen cDNA library or a normalized library containing 12,000 human ORFs fused to the GAL4 activation domain (GAL4-AD; prey libraries). Yeast diploids were grown on appropriate medium for the selection of bait-prey interactions, and positive colonies were analyzed by PCR and sequencing for identifying human proteins captured by N^mono^ in the Y2H system. This screen allowed us to identify, among others, the protein TAX1BP1 as an interactor of the N^mono^ protein (Table 1). For this specific interaction, 40 positive yeast colonies were obtained, and the alignments of the reads from the PCR products showed that the C-terminal part of TAX1BP1 (residues 401-789) is involved in the interaction with N (Figure 1A). None of the cDNA clones expressed full-length TAX1BP1. This probably reflects the fact that isolated domains often better perform than full length proteins in the Y2H system as the reconstitution of a functional GAL4 transcription factor is usually facilitated (42).

**Table 1.**
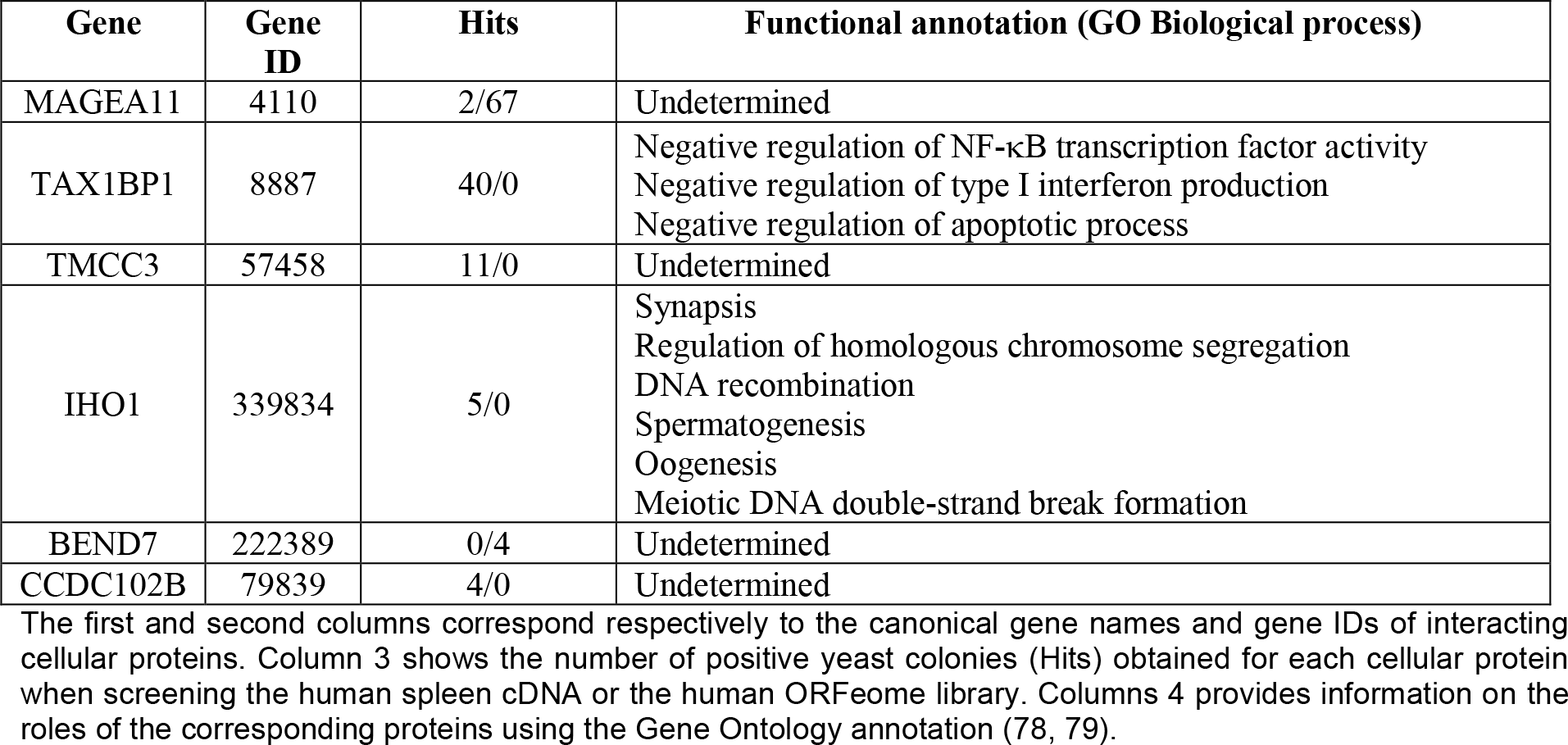
Cellular proteins interacting with RSV N^mono^ identified by Y2H screening.

| Gene | Gene ID | Hits | Functional annotation (GO Biological process) |
| --- | --- | --- | --- |
| MAGEA11 | 4110 | 2/67 | Undetermined |
| TAX1BP1 | 8887 | 40/0 | Negative regulation of NF-κB transcription factor activity<br>Negative regulation of type I interferon production<br>Negative regulation of apoptotic process |
| TMCC3 | 57458 | 11/0 | Undetermined |
| IHO1 | 339834 | 5/0 | Synapsis<br>Regulation of homologous chromosome segregation<br>DNA recombination<br>Spermatogenesis<br>Oogenesis<br>Meiotic DNA double-strand break formation |
| BEND7 | 222389 | 0/4 | Undetermined |
| CCDC102B | 79839 | 4/0 | Undetermined |

The first and second columns correspond respectively to the canonical gene names and gene IDs of interacting cellular proteins. Column 3 shows the number of positive yeast colonies (Hits) obtained for each cellular protein when screening the human spleen cDNA or the human ORFeome library. Columns 4 provides information on the roles of the corresponding proteins using the Gene Ontology annotation (78, 79).

**Figure 1:**
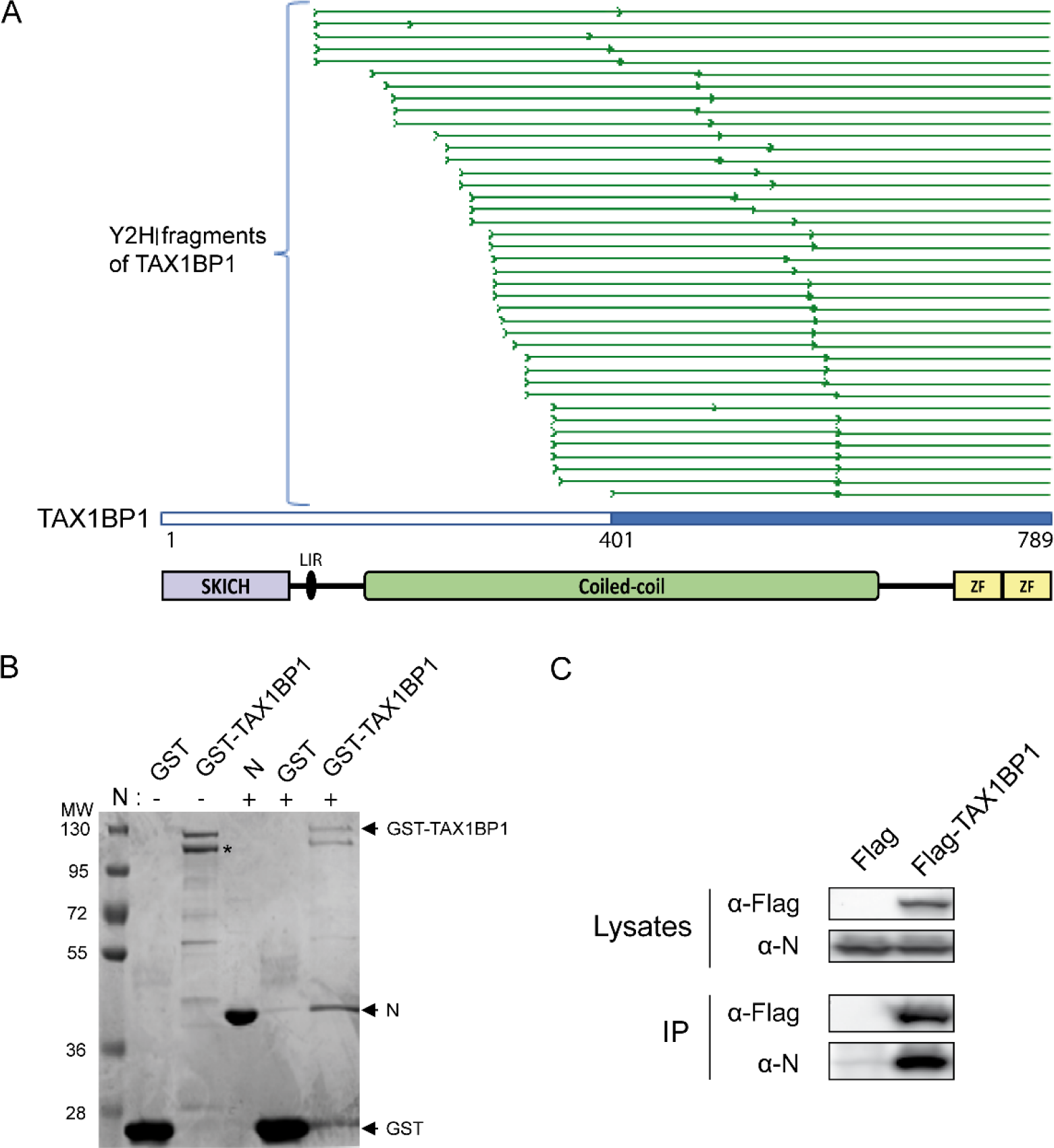
Identification and validation of TAX1BP1-N interaction. **(A)** Multiple alignment of sequencing reads obtained from the 40 yeast colonies matching TAXBP1. As the cDNA library used in the screen was built by oligo-dT priming, TAX1BP1 fragments captured in the screen extend from the beginning of the sequencing reads (thick green line) to the end of the TAX1BP1 sequence. The shortest TAX1BP1 fragment captured with N^mono^ is depicted in blue. Bellow the alignment, a scheme of TAX1BP1 structural organization is presented: SKIP carboxyl homology domain (SKICH), LC3-interacting region (LIR), central coiled coils constituting the oligomerization domain, and the two C-terminal zinc fingers (ZF). **(B)** Validation of N-TAX1BP1 interaction by GST-pulldown with recombinant proteins. GST and GST-TAX1BP1 proteins were purified on glutathione-Sepharose beads and incubated in the presence of recombinant N protein, and interactions was analyzed by SDS-PAGE and Coomassie blue staining. The asterisks indicate the product of degradation of GST-TAX1BP1 corresponding to the deletion of the C-terminal domain. Molecular masses (MW) corresponding to the ladder’s bands are indicated. **(C)** Western blot analysis of the TAX1BP1-N interaction after immunoprecipitation assay. Cells were transiently transfected with constructs allowing the expression of Flag tag alone or the Flag-TAX1BP1 fusion protein with N protein. Immunoprecipitations (IP) were performed with an anti-Flag antibody.

To validate the interaction between TAX1BP1 and the RSV-N protein, we then performed pulldown assays using recombinant proteins. Analysis of purified GST-TAX1BP1 by SDS-PAGE stained with Coomassie blue revealed two main bands with apparent MW close to 120 kDa (Figure 1B). Mass spectrometry analysis of these products allowed to identify the higher migrating band as full length GST-TAX1BP1 (theorical mass,112 kDa), whereas the lower band corresponds to GST-TAX1BP1 partially degraded at the C-terminus (data not shown). When co-incubated with Sepharose-glutathione beads bound to either GST or GST-TAX1BP1, recombinant N protein was specifically captured in the presence of GST-TAX1BP1 (Figure 1B). This result confirmed that RSV-N and TAX1BP1 can directly interact. Finally, we investigated the capacity of RSV-N protein to interact with TAX1BP1 in mammalian cells. Cells were co-transfected with plasmids encoding RSV-N and Flag-tagged TAX1BP1 or the Flag-tag alone as a control, and an immunoprecipitation assay was performed using an anti-Flag antibody. As shown on figure 1C, the RSV-N protein co-precipitated specifically with Flag-TAX1BP1. Altogether, our results indicate that the RSV-N protein can interact directly with TAX1BP1.

### Downregulation of TAX1BP1 expression has limited impact on RSV replication in human cells

TAX1BP1 was recently shown to control the cellular antiviral response during RSV infection (43). We thus determined whether downregulation of TAX1BP1 expressionhas an impact on RSV replication in cell culture (44). Human epithelial A549 cells were transfected with control siRNA (siCT) or siRNA targeting TAX1BP1 (siTAX1BP1). After 24 h of culture, cells were infected with recombinant strains of human RSV expressing either the fluorescent protein mCherry (rHRSV-mCherry) or the bioluminescent enzyme firefly luciferase (rHRSV-Luc). After 48h of culture, mCherry and luciferase expression were determined as a proxy for viral infection. Lower signals were observed in siTAX1BP1-treated cells, thus suggesting a role of TAX1BP1 in RSV replication (Figure 2A). Western-blot analysis of cell lysates confirmed that TAX1BP1 expression is suppressed at this time point (Figure 2B). Somewhat unexpectedly, RSV-N expression in siTAX1BP1-treated cells was similar to control cells (Figure 2B), suggesting that TAX1BP1 has no impact on viral replication. We then further assessed the impact of TAX1BP1 downregulation on viral shedding by quantifying virions in culture supernatants of infected cells. As shown on figure 2C, viral titers in supernatants of siTAX1BP1-treated cells were similar to matching siCT-treated controls. Altogether, these results led to the conclusion that TAX1BP1 is not directly involved in RSV replication. We therefore decided to explore a more indirect role of TAX1BP1 on RSV replication that would depend on its regulatory role on the innate immune response.

**Figure 2:**
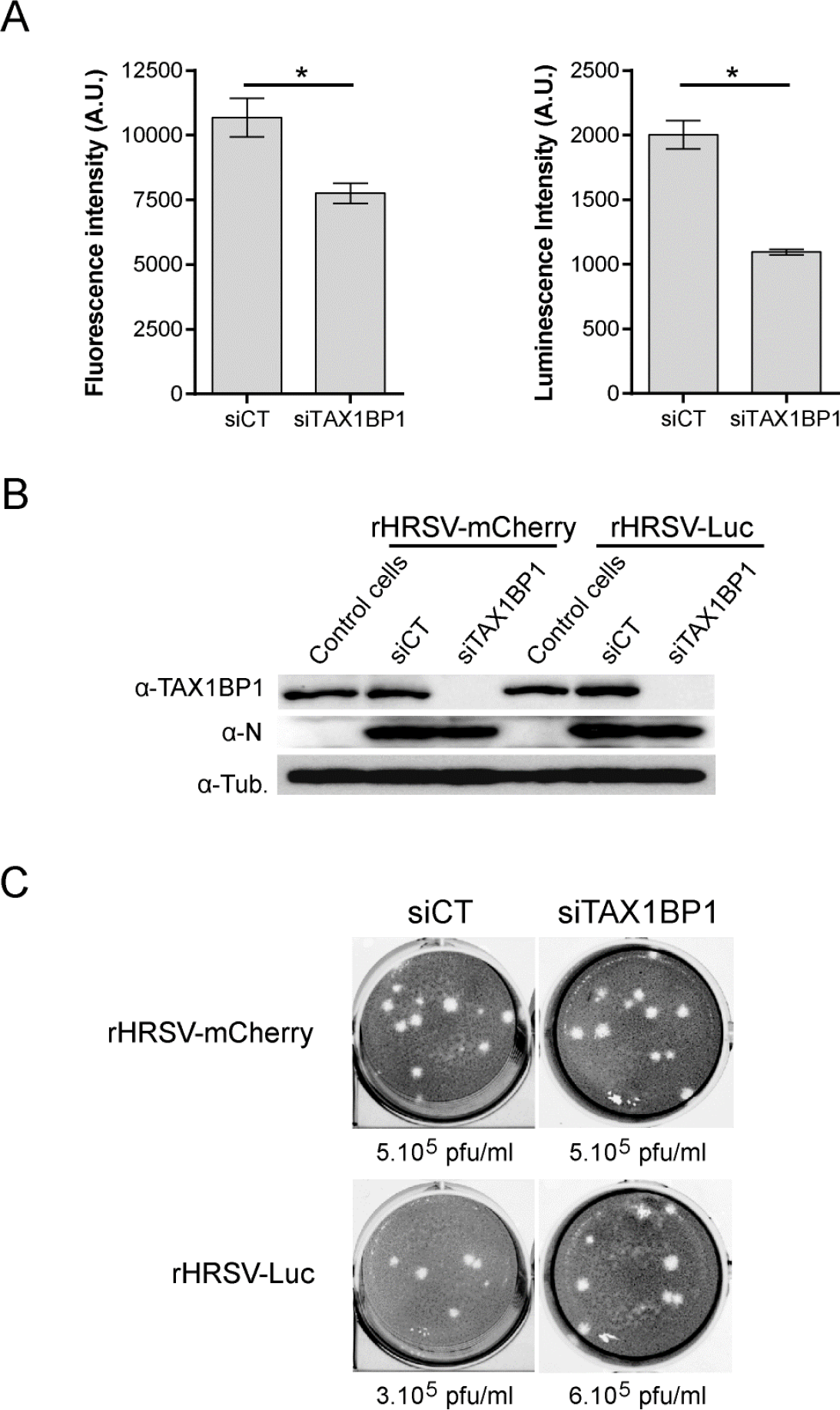
Impact of TAX1BP1 depletion on RSV replication in cells. A549 cells were transfected with siRNAs control (siCT) or targeting TAX1BP1 (siTAX1BP1) and then infected 24 h later with either rHRSV-mCherry or rHRSV-Luc, at a MOI of 0.5. **(A)** RSV replication was quantified 48 h post-infection by measurement of fluorescence (left) and luminescence (right) expressed in arbitrary unit (A.U.) in cell lysates. Data are representative of three experiments made in quadruplicates. Data are mean ± SEM, *p < 0.05. **(B)** Western blot analysis of TAX1BP1 silencing and RSV N expression in cells infected with either rHRSV-mCherry or rHRSV-Luc, 48 h post-infection. (C) Titration of virions released in the culture media of cells treated with siCT (left) and siTAX1BP1 (right) and infected with rHRSV-mCherry (upper panel) or rHRSV-Luc (lower panel). Calculated viral titers in plaque-forming unit per ml (pfu/ml) are indicated.

### Depletion of TAX1BP1 impairs RSV replication in mice

Given the complexity of the immune response triggered upon RSV infection, we assessed the impact of TAX1BP1 depletion directly *in vivo* using TAX1BP1-deficient (TAX1BP1^KO^) mice. These mice being generated in 129-strain mice (39), we first investigated the kinetics of rHRSV-Luc replication in this genetic background. Although luminescence was shown to be correlated to viral replication by direct measurement on live animals in BALB/c mice using the IVIS system (44, 45), the skin pigmentation of 129 mice impaired luminescence detection. We thus decided to monitor viral replication in infected animals by measuring the luciferase activity in lung homogenates. Wild-type 129 mice were either instilled with mock control (Mock) consisting of HEp2 cell culture supernatant, or infected with 1.87 x 10^5^ pfu of rHRSV-Luc *via* intranasal (IN) inoculation. The viral replication was quantified the first 4 days post-infection (p.i.). The bioluminescence in lung homogenates was detected at day 1 p.i., and viral replication in the lungs increased from day 2 to day 4 p.i. (Figure 3A, left). In parallel, expression of *N-RSV* gene in the lung lysates was quantified by qRT-PCR (Figure 3A, right). Data showed that N mRNA could be detected from day 2 p.i., and that the peak of infection was reached at day 3 and 4 p.i.. These results revealed a respectable correlation between bioluminescence intensity and RSV N mRNA expression in line with previous reports (44), with a clear detection of RSV replication at day 3 and 4 p.i.. Of note, this kinetics of replication is similar to the one described in BALB/c mice, a reference mouse strain to study RSV infection (44, 45).

**Figure 3:**
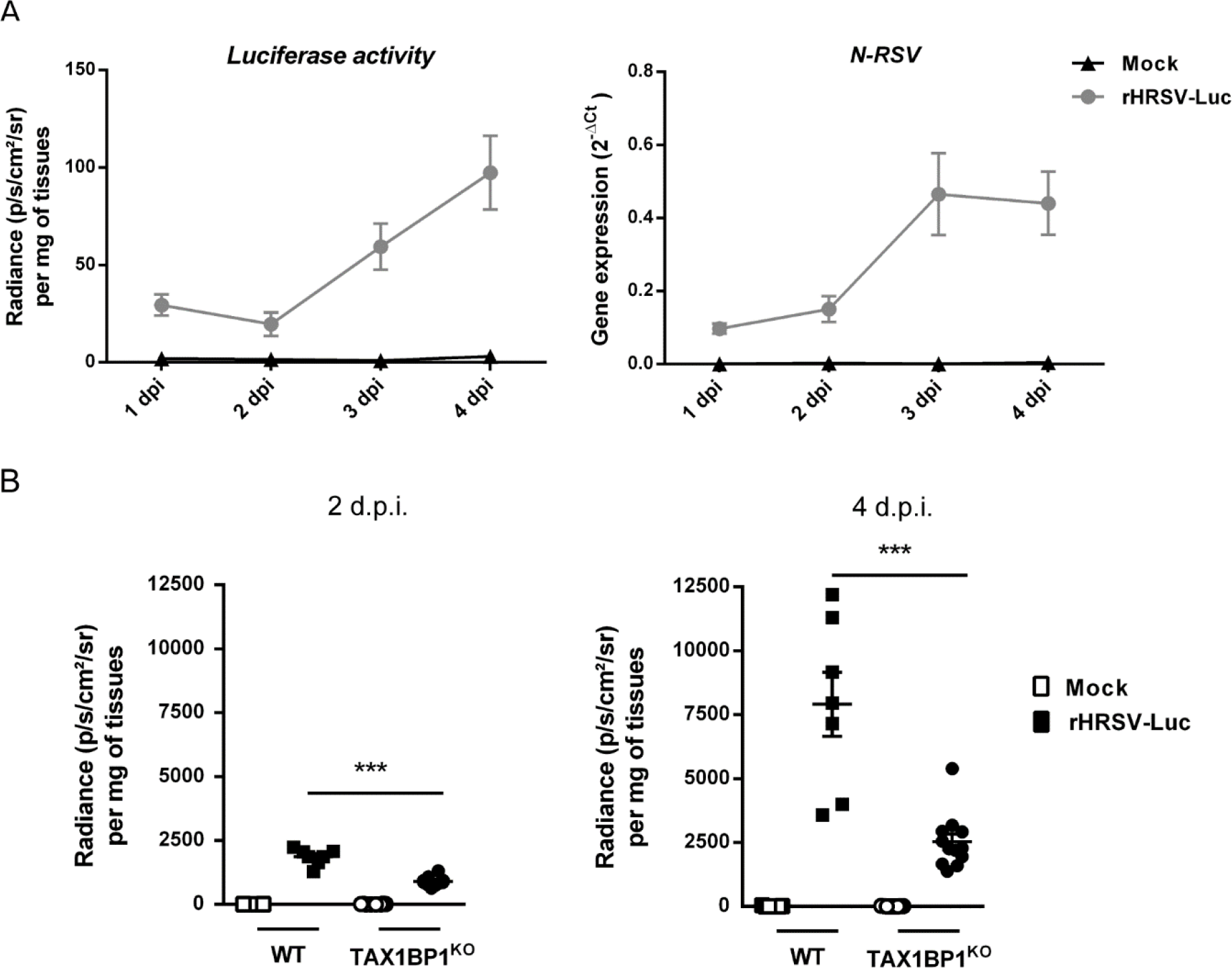
TAX1BP1-deficient mice infected with RSV present a reduced virus replication in the lungs. **(A)** Kinetics of RSV infection in 129 mice. Wild-type (WT) strain 129 mice were infected with Hep2-supernatant (Mock, n = 1) or rHRSV-Luc (n = 4). (Left) Luciferase activity associated to viral replication was measured at different days post-infection (d.p.i.) in lung lysates, by quantification of photon emission (radiance in photon/sec/cm^2^/sr) and normalized to the amount of lysed tissue. (Right) In parallel, *N-RSV* gene expression was measured in the lung lysates by RT-qPCR and calculated by the formula 2^-ΔCt^ with ΔCT = Ct_N-RSV_ - Ct_HPRT_. Data are mean ± SEM, *p < 0.05. **(B)** WT or TAX1BP1^KO^ 129 mice were infected with HEp2-supernatant (Mock) or rHRSV-Luc. Luciferase activity associated to viral replication was measured at 2 or 4 d.p.i. (left and right respectively) in lung lysates, by quantification of photon emission (radiance in photon/sec/cm^2^/sr) and normalized to the amount of lysed tissue. Data are mean ± SEM from two independent experiments with *n* = 7 for RSV infected WT mice and *n* = 11 for RSV infected TAX1BP1^KO^ mice.

Based on these results, we decided to compare rHRSV-Luc replication in wild-type (WT) and TAX1BP1^KO^ 129 mice. We chose to quantify bioluminescence in the lung of mock-treated and HRS-infected animals at day 2 and day 4 p.i. in order to compare viral replication at an early time point and at the peak of infection. Of note, it was previously established that viral replication estimated *in vivo* by quantifying bioluminescence signals directly correlates with viral load (44). Our results showed a strong reduction in RSV replication in TAX1BP1^KO^ mice compared to WT mice, at both day 2 p.i. and 4 p.i. (Figure 3B). These results revealed a critical role of TAX1BP1 on RSV replication *in vivo*.

### Depletion of TAX1BP1 favors antiviral and inflammatory responses during RSV infection

As mentioned, among the various functions of TAX1BP1, this cellular protein acts as a cofactor of the A20 protein, which is a negative regulator of NF-κB and IRF3/7 pathways that are respectively involved in inflammatory and antiviral responses. In the mouse model of RSV infection, the induction of inflammatory cytokines and IFN-I in the first hours post-exposure to the virus are well documented (46-49). We thus assessed if the inhibition of RSV replication upon TAX1BP1 depletion could be associated to a modulation of the antiviral and inflammatory responses in the lungs of infected mice at early time point post-infection. Mice were mock-treated or infected with 1.87 x 10^5^ pfu of rHRSV-Luc and at day 1 p.i., expression levels of IFN-I (IFN-α and IFN-β) and of the inflammatory cytokines IL-6 and TNF-α were determined from lung lysates of WT or TAX1BP1^KO^ mice. As shown on Figure 4, RSV infection induced the production of IFN-α, IFN-β, IL-6 and TNF-α in all the animals. Slightly but significantly higher levels of IFN-α and TNF-α were detected in the lungs of TAX1BP1^KO^ mice compared to WT mice (Figure 4A and D). However, no difference in IFN-β or IL-6 expression was detected between infected WT and TAX1BP1^KO^ mice (Figure 4B and C). Of note, all groups of animals showed comparable levels of RSV infection as assessed by bioluminescence quantification in the lung homogenates (not shown). Because the measurements were performed in whole lung lysates, the quantified cytokines and chemokines are probably produced by several cell populations (*i*.*e*. both epithelial and immune cells). We thus decided to specifically focus on alveolar macrophages (AMs) which are main actors of the antiviral response to RSV (47). AMs were isolated from WT and TAX1BP1^KO^ mice after repeated bronchoalveolar lavages and cultured for 24 h before incubation for another 24 h in the presence of either rHRSV-mCherry or UV-inactivated rHRSV-mCherry (MOI=5). Culture supernatants were collected, and IFN-α, IFN-β, IL-6 and TNF-α were quantified by immunoassay. A strong induction of both anti-viral (Figure 5A and B) and inflammatory cytokines (Figure 5C and D) was detected in the supernatant of AMs exposed to RSV, whereas a much weaker induction of these molecules was observed for AMs exposed to inactivated RSV, thus validating an efficient infection of AMs. Of note, although AMs can be infected by RSV, these cells do not productively replicate the virus (50). Most interestingly, the production of IFN-α, IFN-β, IL-6 and TNF-α was enhanced in AMs derived from TAX1BP1^KO^ mice compared to AMs isolated from WT mice (Figure 5). Altogether, these results demonstrate that TAX1BPA1 is a key factor involved in the inhibition of the antiviral and inflammatory responses in the lungs of RSV-infected animals and in isolated AMs.

**Figure 4:**
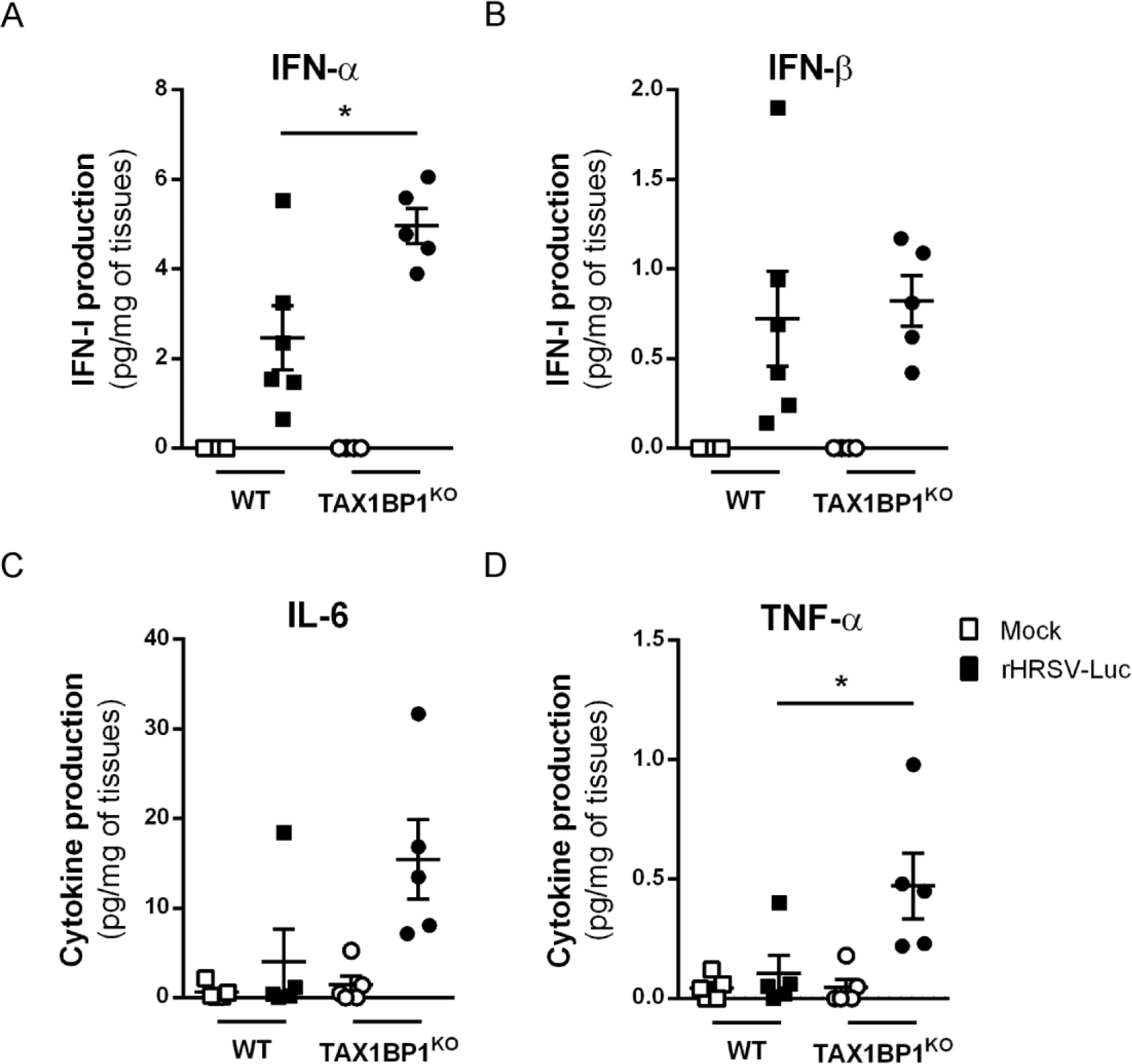
Study of antiviral/inflammatory immune responses in the lungs of infected TAX1BP1^KO^ mice. WT or TAX1BP1^KO^ mice were infected with HEp2-supernatant (Mock) or rHRSV-Luc. (**A, B**) The productions of IFN-α and IFN-β were measured 24 h post-infection in lung lysates using ProcartaxPlex immunoassay. (**C, D**) The productions of IL-6 and TNF-α were measured 24 h post-infection in lung lysates using MilliPlex MAP immunoassay. The concentrations were normalized to weight lungs. Data are mean ± SEM and are representative of two independent experiments with *n* = 5-6 mice per group.

**Figure 5:**
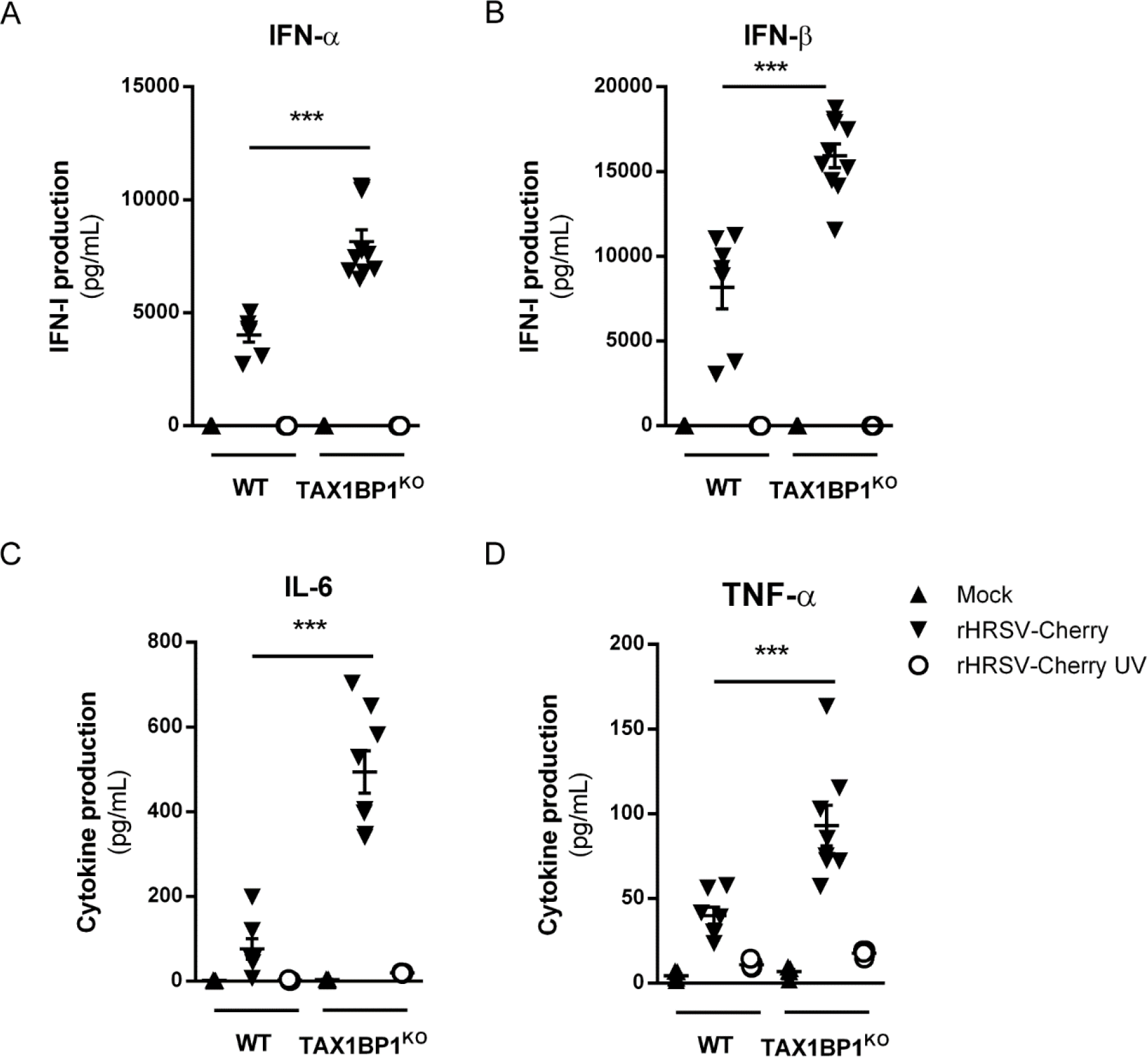
Deletion of TAX1BP1 enhances the production of type I IFN and inflammatory cytokines in AMs following RSV infection. AMs from WT or TAX1BP1^KO^ mice were either not infected (mock, black triangle) or exposed to rHRSV-mCherry (RSV, inverted black triangle symbol) or UV-inactivated rHRSV-mCherry (UV-RSV, white circle) at MOI of 5 for 2h. (**A, B**) The productions of IFN-α and IFN-β were measured 24h post-infection in supernatants using ProcartaxPlex immunoassay. (**C, D**) The productions of IL-6 and TNF-α were measured 24 h post-infection in supernatants using MilliPlex MAP immunoassay. Data are mean ± SEM from two independent experiments.

## DISCUSSION

Previous studies using microarray and proteomic approaches have provided key information on RSV-host interactions (51, 52), but the interactome of RSV proteins still remains poorly characterized. Due to their pivotal role during virus entry, replication and assembly, it is expected that components of the viral polymerase complex, and especially the N protein, are involved in various interactions with cellular factors. The objective of this study was to find new cellular partners of RSV-N by performing a yeast two-hybrid screen. Using this approach, we captured 6 cellular proteins using RSV-N as bait, among which TAX1BP1 was overrepresented. We thus focused on TAX1BP1 as TAX1BP1 depletion has recently been shown to favor the innate immune response to RSV infection and to impair viral replication in cell culture (43). In addition, TAX1BP1 is already known to interact with different viral proteins including the N protein of measles virus that belongs to *Mononegavirales* order like RSV (35-38), suggesting that this protein is often hijacked by viruses. TAX1BP1 is a homodimer of about 90 kDa and is organized into three main structural domains. The N-terminal SKIP carboxyl homology (SKICH) domain (53) was recently shown to interact with the adaptor protein NAP1, allowing the recruitment of the TANK-binding kinase 1 (TBK1), which is involved in selective autophagy of invading pathogens and damaged mitochondria but is also critical to the induction of IFN-I by RIG-I, MDA5 and STING (54-58). It is followed by a LC3-interacting region (LIR) that can bind different LC3/GABARAP orthologs (59) involved in the recruitment of TAX1BP1 to autophagosomes. The central part of TAX1BP1 exhibits coiled coils forming the oligomerization domain that interacts with TRAF6 protein (60), and is followed by two C-terminal zinc fingers (UBZ1 and UBZ2) (61). These zing fingers were shown to interact with ubiquitinylated proteins, with myosin VI, and with the protein A20 (62-64).

Here, the alignment of the PCR reads obtained from the 40 yeast clones that expressed TAX1BP1 in the two-hybrid screen revealed that the C-terminal part of this protein is involved in the interaction with RSV-N. Based on our results, it is expected that the TAX1BP1 binding site to RSV-N is located within the oligomerization domain and/or the C-terminal zinc finger domains. The N-TAX1BP1 interaction was validated first by pulldown using recombinant TAX1BP1 and RSV-N proteins, and then by immunoprecipitation when co-expressing the two proteins in human cells. Noteworthy, we managed to purify the recombinant TAX1BP1 protein to validate the direct interaction with the RSV-N protein. However, the purification of this protein was challenging as TAX1BP1 tends to be cleaved at its C-terminus, and this hampered affinity study with RSV-N by biophysical approaches. To gain structural and functional insights on this interaction that could represent a new therapeutic target, a precise characterization of TAXBP1 binding domains to RSV-N is required. The structure of the C-terminal UBZ domain of TAXBP1 either alone or in complex with Myosin VI has already been resolved (61, 64). The crystal structure of RSV nucleocapsid-like structures consisting of rings containing 10 N protomers and RNA of 70 nucleotides has been determined (65). Recently, a recombinant RSV N^0^-P complex has also been characterized (66). The reconstitution of a recombinant complex of RSV-N (monomeric or oligomeric form) bound to the C-terminal fragment of TAX1BP1 could thus provide key structural information on this interaction. Finally, given the strong homology between the N proteins of RSV and human Metapneumovirus (hMPV), another pneumovirus also responsible of acute respiratory infections, the potential interaction between hMPV-N and TAX1BP1, and its functional relevance during infection should also be investigated.

We then investigated the potential role of TAX1BP1 in RSV infection. TAX1BP1 suppression showed a limited or no impact on viral protein expression in cell culture, and the production of new viral particles was unaffected. However, a model of RSV-infected TAX1BP1^KO^ mice unraveled the critical role of TAX1BP1 in RSV infection *in vivo*, the depletion of TAX1BP1 leading to a nearly 3-fold decrease in viral replication in the lungs of infected mice. We also showed that RSV-infected TAX1BP1^KO^ mice present higher levels of IFN-α and TNF-α in the lungs compared to WT mice at day 1 p.i. Besides, RSV-infected AMs isolated from TAX1BP1^KO^ mice produced higher levels of IFN-I (IFN-α and β) and inflammatory cytokines (IL-6 and TNF-α) compared to those isolated from WT mice.

These results reveal that TAX1BP1 participates to the attenuation of the host antiviral and inflammatory responses during RSV infection *in vivo* and especially in AMs. Altogether, this suggests that TAXBP1 recruitment by RSV-N indirectly promotes RSV growth by inhibiting the innate immune response. Overall, this conclusion is consistent with the recent study by Martín-Vicente *et al*. (43) but significant differences should be highlighted. Indeed, they found that the production of infectious RSV particles in A549 cells decreases when silencing TAX1BP1 or interacting co-factors A20, ABIN1 and ITCH. In our hands, the effect of TAX1BP1 silencing on RSV infection was striking only *in vivo*. At this point, we don’t have an explanation to this discrepancy as we be both used the same *in vitro* model of A549-infected cells. Besides, they found in their study that A549 cells silenced for TAX1BP1 express higher levels of ISG15, IL-6 and IL-8 upon RSV infection, but IFN-β and TNF-α expression were not significantly affected. On the contrary, we found that TAX1BP1-deficient AMs express higher level of TNF-α, IL6, IFN-β and IFN-α when infected by RSV. The use of distinct cellular models and TAX1BP1-depletion methods could account for these differences. Indeed, TAX1BP1 is directly involved in the regulation of innate immune pathways, but is also an adaptor for autophagy (62) which is required for the induction of an optimal antiviral response in RSV-infected macrophages (67). Thus, the role of TAX1BP1 in the regulation of the innate immune response induced upon RSV infection could vary between epithelial and immune cells depending on the relative contribution of autophagy in the activation of the innate immune response. Finally, it should be noticed that TAX1BP1 has been previously described to regulate B cell differentiation (68). It would thus be interesting to study whether TAX1BP1 could also be involved in acquired immune responses in the context of RSV infection *in vivo*, and in particular the production of antibodies.

As TAX1BP1 works an adaptor protein in different processes, it is essential to characterize TAX1BP1 partners in different cell lines when infected by RSV. During our study, we investigated the cellular localization of TAX1BP1 in the context of viral infection or overexpression of N, in order to determine in particular if TAX1BP1 could be recruited to IBs, as previously shown for MDA5 and MAVS (32), or if TAXBP1 could recruit RSV-N to specific cellular compartments. However, we didn’t manage to clearly detect TAX1BP1 by immunolabeling using commercial antibodies. Furthermore, upon overexpression of Flag- or GFP-tagged TAX1BP1 in cells, TAX1BP1 was shown to concentrate into cytoplasmic granules and to induce cell death, thus precluding further analysis (data not shown).

In conclusion, we have shown that TAX1BP1 is suppressing the innate immune response to RSV *in vivo* and in AMs. Results also suggest that RSV hijacks this mechanism through a direct physical interaction with RSV-N. Although the precise role of TAX1BP1 in RSV infection needs to be further characterized, this interaction helps understanding the pathogeny associated to the infection and represents new target for antiviral approaches.

## MATERIALS AND METHODS

### Plasmids and siRNA

The plasmid pFlag-TAX1BP1 encoding for TAX1BP1 in fusion with a N-terminal Flag tag was kindly provided by Dr C. Journo (ENS, Lyon, France). The plasmid pFlag was obtained by inserting a stop codon in the pFlag-TAX1BP1 vector, using the Quickchange site-directed mutagenesis kit (Stratagene). The already described p-N (69) was used for cell transfection and immunoprecipitation assay.

The pGEX-4T-3 vector was used to produce recombinant Glutathione S-transferase protein (GST). The pGEX-TAX1BP1 plasmid expressing the GST in fusion with the N-terminus of TAX1BP1 was obtained by cloning the TAX1BP1 sequence between BamHI and XhoI sites of the pGEX-4T-3 plasmid. For purification of recombinant N protein, the pET-N and pGEX-PCT plasmids already described (41) were used. For yeast two-hybrid screening, the DNA sequence encoding the N^mono^ (monomeric N mutant K170A/R185A) was cloned by *in vitro* recombination (Gateway technology; Invitrogen) from pDONR207 into the yeast two-hybrid vector pPC97-GW for expression in fusion downstream of the GAL4 DNA-binding domain (GAL4-BD). The control siRNA and a pool of TAX1BP1 siRNA (Ambion) were used for TAX1BP1 silencing experiments.

### Antibodies

The following primary antibodies were used for immunoprecipitation assay and/or immunoblotting: a mouse anti-Flag and a mouse anti-Flag-HRP antibody (Sigma), a rabbit anti-N antiserum (70), and a mouse monoclonal anti-β-tubulin antibody (Sigma). Secondary antibodies directed against mouse and rabbit Ig G coupled to HRP (P.A.R.I.S) were used for immunoblotting.

### Cell lines

BHK-21 cells (clone BSRT7/5), hamster kidney cells constitutively expressing the T7 RNA polymerase (71), HEp-2 cells (ATCC number CCL-23), and human lung carcinoma epithelial A549 cells were grown in Dulbeco Modified Essential Medium (Lonza) supplemented with 10% fetal calf serum (FCS), 2 mM glutamine, and 1% penicillin-streptomycin. The transformed human bronchial epithelial cell line BEAS-2B (ATCC) was maintained in RPMI 1640 medium (Invitrogen) supplemented with 10% fetal bovine serum (FBS, Invitrogen), 1% L-glutamine, and 1% penicillin-streptomycin.

### Viruses

Recombinant RSV viruses rHRSV-mCherry and rHRSV-Luc corresponding to RSV Long strain expressing either the mCherry or the Luciferase proteins were amplified on HEp-2 cells and titrated using a plaque assay procedure as previously described (44). Briefly for titration cells were infected with serial 10-fold dilutions of viral supernatent in complete minimum essential medium (MEM). The overlay was prepared with microcrystalline cellulose Avicel RC581 (FMC Biopolymer) at a final concentration of 0.6% in complete MEM containing 1% fetal calf serum. After 6 days at 37°C and 5% CO2, plaques were revealed by 0.5% crystal violet / 20% ethanol solution staining of the cell layers, and the number of plaque-forming unit (pfu) per well was counted.

### Yeast Two-Hybrid Screening

Yeast two-hybrid screens were performed following the protocol described in Vidalain et al. (72). AH109 yeast cells (Clontech; Takara, Mountain View, CA, USA) were transformed with pGAL4-BD-N^mono^ using a standard lithium-acetate protocol. Screens were performed on a synthetic medium lacking histidine (-His) and supplemented with 3-amino-1,2,4-triazole (3-AT) at 10 mM. A mating strategy was used to screen two different prey libraries with distinct characteristics: a human spleen cDNA library, and a normalized library containing 12,000 human ORFs (73). All libraries were established in the yeast two-hybrid expression plasmid pPC86 to express prey proteins in fusion downstream of the GAL4 transactivation domain (GAL4-AD). After six days of culture, colonies were picked, replica plated, and incubated over three weeks on selective medium to eliminate potential contamination with false positives. Prey proteins from selected yeast colonies were identified by PCR amplification using primers that hybridize within the pPC86 regions flanking the cDNA inserts. PCR products were sequenced, and cellular interactors were identified by multi-parallel BLAST analysis.

### Expression and purification of recombinant proteins

*E. coli* BL21 bacteria (DE3) (Novagen, Madison, WI) transformed with pGEX-4T-3 and pGEX-TAX1BP1 plasmids were grown at 37°C for 2-3 h in 200 mL of Luria Bertani (LB) medium containing 100 µg/mL ampicillin until the OD_600nm_ reached 0.6. Protein expression was then induced by addition of 1 mM of isopropyl-ß-D-thio-galactoside (IPTG) in the presence of 50 mM ZnSO_4_ during 4 h at 37°C before harvesting by centrifugation. Expression and purification of the recombinant N protein was previously described (65, 74). Briefly, BL21 bacteria co-transformed with pET-N-pGEX-PCT plasmids were grown in LB medium containing kanamycin (50 µg/mL) and ampicillin for 8h at 37°C. Then, the same volume of fresh LB was added and protein expression was induced by adding IPTG at 80 µg/ml to the culture. The bacteria were incubated for 15 h at 28°C and then harvested by centrifugation. For GST-fusion proteins purification, bacterial pellets were re-suspended in lysis buffer (50 mM Tris-HCl pH 7.8, 60 mM NaCl, 1 mM EDTA, 2 mM DTT, 0.2% Triton X-100, 1 mg/mL lysozyme) supplemented with complete protease inhibitor cocktail (Roche, Mannheim, Germany), incubated for 1 hour on ice, sonicated, and centrifuged at 4°C for 30 min at 10,000 g. Glutathione-Sepharose 4B beads (GE Healthcare, Uppsala, Sweden) were added to clarified supernatants and incubated at 4°C for 15 h. Beads were then washed two times in lysis buffer and three times in PBS 1X, then stored at 4°C in an equal volume of PBS. To isolate the recombinant N protein, beads containing bound GST-PCT+N complex were incubated with thrombin (Novagen) for 16 h at 20°C. Purified recombinant N proteins were loaded onto a Superdex 200 16/30 column (GE Healthcare) and eluted in 20 mM Tris/HCl pH 8.5, 150 mM NaCl.

#### Pulldown assays

Purified recombinant N protein was incubated in the presence of GST or the GST-TAX1BP1 fusion protein fixed on beads in a final volume of 100 µL in buffer Tris 20 mM, pH 8.5, NaCl 150 mM. After 1 h under agitation at 4°C, the beads were extensively washed with 20 mM Tris (pH 8.5)–150 mM NaCl, boiled in 30 µL Laemmli buffer, and analyzed by SDS-PAGE and Coomassie blue staining.

#### Coimmunoprecipitation assay

BSRT-7 cells were cotransfected with pFlag or pFlag-TAX1BP1 and pN for 36 h. Transfected cells were then lysed for 30 min at 4°C in ice-cold lysis buffer (Tris HCl 50 mM, pH 7.4, EDTA 2 mM, NaCl 150 mM, 0.5% NP-40) with a complete protease inhibitor cocktail (Roche), and coimmunoprecipitation experiments were performed on cytosolic extracts. Cell lysates were incubated for 4 h at 4°C with an anti-Flag antibody coupled to agarose beads (Euromedex). The beads were then washed 3 times with lysis buffer and 1 time with PBS, and proteins were eluted in Laemmli buffer at 95°C for 5 min and then subjected to SDS-PAGE and immunoblotting.

### siRNA transfection and infection

Freshly passaged A549 cells were transfected with the indicated siRNA at a final concentration of 10 nM by reverse transfection into 48 wells plates, using Lipofectamine RNAiMAX (ThermoFischer) according to the manufacturer’s instructions. Briefly, a mixture containing Opti-MEM (Invitrogen), lipofectamine RNAiMAX and siRNA was incubated for 5 min at room temperature before depositing at the bottom of the wells. The cells in DMEM medium without antibiotics were then added dropwise before incubation at 37°C, 5% CO_2_. After 24 h of transfection in the presence of siRNA, the medium was removed and the cells were infected with recombinant rHRSV-mCherry or rHRSV-Luc viruses at a MOI of 0.5 in DMEM medium without phenol red and without SVF, for 2 h at 37°C. The medium was then replaced by DMEM supplemented with 2% SVF and the cells were incubated for 48 h at 37°C. For cells infected with the rHRSV-mCherry virus, the quantification of replication was performed by measuring the mCherry fluorescence (excitation: 580 nm, emission: 620 nm) using a Tecan Infinite M200 Pro luminometer. For HRSV-Luc replication quantification, cells were lysed in luciferase lysis buffer (30 mM Tris pH 7.9, 10 mM MgCl2, 1 mM DTT, 1% Triton X-100, and 15% glycerol). After addition of luciferase assay reagent (Promega), luminescence was measured using a Tecan Infinite M200 Pro luminometer. Non-infected A549 cells were used as standards for fluorescence or luminescence background levels. Each experiment was performed in triplicates and repeated at least three times. For each experiment, cells treated in the same conditions were lysed and protein expression was analyzed by Western blotting.

### RSV infection of mice and luciferase measurement

TAX1BP1-deficient (TAX1BP1^KO^) 129 mice were created by gene targeting, as previously described (39). TAX1BP1^KO^ mice and wild-type 129 co-housed control animals were bred and housed under SPF conditions in our animal facilities (IERP, INRAE, Jouy-en-Josas). Wild type (WT) and TAX1BP1^KO^ female and male mice at 8 weeks of age (n=11 per group) were anesthetized with of a mixture of ketamine and xylazine (1 and 0.2 mg per mouse, respectively) and infected by intranasal administration of 80 μL of recombinant RSV expressing luciferase (rHRSV-Luc, 2.34 x 10^6^ pfu/mL) (44, 75, 76) or cell culture media as mock-infection control. Mice were then sacrificed at different timepoints by intraperitoneal (I.P.) injection of pentobarbital and lungs were frozen.

### Viral *N-RNA* gene expression by RT-qPCR

Frozen lungs were homogenized in NucleoSpin®RNA XS Kit (Macherey-Nagel) lysis buffer with a Precellys 24 bead grinder homogenizer (Bertin Technologies, St Quentin en Yvelines, France). Total RNA was extracted from lungs or infected cells using NucleoSpin® RNA kit (Macherey-Nagel) and reverse transcribed using the iScript™ Reverse Transcription Supermix for RT-qPCR kit (Bio-Rad) according to the manufacturer’s instructions. The primers (Sigma-Aldrich) used are listed below. The qPCRs were performed with the MasterCycler RealPlex (Eppendorf) and SYBRGreen PCR Master Mix (Eurogenetec) and data analyzed with the Realplex software (Eppendorf) to determine the cycle threshold (Ct) values. Results were determined with the formula 2^-ΔCt^ with ΔCT = Ct_gene_-Ct_HPRT._ The primers (Sigma-Aldrich) used are listed below: HPRT (hypoxanthine-guanine phosphoribosyltransferase), Forward primer 5’-CAGGCCAGACTTTGTTGGAT-3’ and Reverse primer 5’-TTGCGCTCATCTTAGGCTTT-3’; and N-RSV, Forward primer 5’-AGATCAACTTCTGTCATCCAGCAA-3’ and Reverse primer 5’-TTCTGCACATCATAATTAGGAGTATCAAT-3’.

### Luciferase expression in lung lysates

Frozen lungs were weighed and then homogenized in 300 μL of Passive Lysis Buffer (PLB) (1 mM Tris pH 7.9; 1 mM MgCl2; 1% Triton × 100; 2% glycerol; 1 mM DTT) with a Precellys 24 bead grinder homogenizer (Bertin Technologies, St Quentin en Yvelines, France) and a cycle of 2 × 15 s at 4 m/s. Lung homogenates were clarified by centrifugation 5 min at 2000 g and distributed on microplates (50 µL). Then, 50 µL of luciferase assay reagent (Promega) were added on each well. The detection of firefly luciferase activity was measured by photon emission using an *In Vivo* Imaging System (IVIS-200, Xenogen, Advanced Molecular Vision) and Live Imaging software (version 4.0, Caliper Life Sciences). Data were expressed in radiance (photons/sec/cm^2^/sr) and normalized to weight lungs.

### RSV infection of AMs

A cannula was inserted in trachea from mice and repeated bronchoalveolar lavages (BALs) were made with PBS. AMs were isolated after centrifugations and 1 x 10^5^ AMs were plated in 96-well cell culture plates in RPMI supplemented with L-glutamine 2 mM, FCS 5% and antibiotics for 24 h to allow for adhesion, as previously described (77). AMs were then exposed to rHRSV-mCherry or ultra-violet (UV)-inactivated rHRSV-mCherry (the same batch exposed 20 min to UV) at MOI 5 or Hep2 cell culture supernatant (Mock). After 24 h, supernatants were collected and were frozen for cytokine quantification.

### Cytokine quantification

IFN-α and IFN-β or IL-6 and TNF-α were measured in supernatants of AMs or lung lysates using IFN alpha/IFN beta 2-Plex Mouse ProcartaPlex™ immunoassay (ebiosciences) or Milliplex MAP Mouse™ assay (Merck), respectively. Data were acquired using a MagPix multiplex system (Merck) in order to determine the mean of fluorescent intensities (MFIs) and results were analyzed on Bio-Plex Manager™ software. The concentrations were normalized to lungs weight.

### Ethics statement

The *in vivo* work of is study was carried out in accordance with INRAE guidelines in compliance with European animal welfare regulation. The protocols were approved by the Animal Care and Use Committee at “Centre de Recherche de Jouy-en-Josas” (COMETHEA) under relevant institutional authorization (“Ministère de l’éducation nationale, de l’enseignement supérieur et de la recherche”), under authorization number 2015060414241349_v1 (APAFIS#600). All experimental procedures were performed in a Biosafety level 2 facility.

### Statistical analysis

Nonparametric Mann-Whitney (comparison of two groups, n ≥ 4) was used to compare unpaired values (GraphPad Prism software). Significance is represented: *p < 0.05; **p < 0.01 and ***p < 0.001;

## Acknowledgments

We thank Dr. Sabine Riffault (INRAE, Jouy-en-Josas) for helpful discussion and critical reading of the manuscript. We are grateful to Chloé Journo (ENS-Lyon, France) for providing the pFlag-TAX1BP1 plasmid, Céline Urien (INRAE, Jouy-en-Josas) for mice genotyping, and the Infectiology of fishes and rodent facility (IERP, INRAE, doi: 10.15454/1.5572427140471238E12) to animals’ facilities and for birth management. We thank the Emerg’in platform for access to IVIS200 that was financed by the Region Ile De France (SESAME), and the Plateforme d’Analyse Protéomique de Paris Sud-Ouest (PAPPSO, INRAE) for mass spectrometry analysis. C. Drajac. and Q. Marquant were recipients of a Ph.D. and Post-doctoral fellowship of the Région Ile-de-France (DIM-Malinf and DIM-OneHealth, respectively), A. Peres de Oliveira was recipient of post-doctoral fellowship (CAPES-Brazil 14809-13-3/ CAPES-COFECUB 769-13). This study was supported in part by Grants-in-Aid for scientific research from the Ministry of Education, Culture, Sports, Science, and Technology, Japan to H.Iha, and with the financial support of the French Agence Nationale de la Recherche, specific program ANR Blanc 2013 “Respisyncycell” (ANR-13-IVS3-0007 and FAPESP-Brazil/ANR - BLANC - RESPISYNCELL 2013/50299-2).

## Author contributions

DD, AMV, JFE, POV and MG designed experiments. APO, SM, LG, JF, FB and MG performed molecular and cellular assays. SM, CD, VP, QM, EB, HI and DD performed mice experiments, samples’ treatment and analysis of *in vivo* experiments. APO, FT and POV performed two hybrid screens. MG, DD, POV and JFE wrote the paper. MG edited the manuscript. All authors commented on the manuscript.

